# Methods for *In situ* Quantification of Mitochondrial Morphology In Muscle and Terminal Schwann Cells of Mice

**DOI:** 10.1101/2025.09.29.679202

**Authors:** Aaron B. Morton, Jacob A. Kendra, Brian Glancy, Shadi Golpasandi, Alexandra G. Naman

**Author notes:** **Correspondence:**Aaron B. Morton, PhD Kinesiology and Sport Management Texas A&M University 314 Gibb Gilchrist Building 2929 Research Pkwy College Station, TX 77845.

## Abstract

Mitochondrial dysfunction is well described in many chronic illnesses including musculoskeletal, neurodegenerative, and cardiovascular diseases. Mitochondrial network morphology has been implicated as a biomarker of disease, correlating increased mitochondrial fragmentation to impaired cellular function. While advancements in imaging techniques further our understanding of mitochondrial dynamics in live cells, easily accessible approaches for accurate quantification of *in situ* mitochondrial networks in low abundance tissues are lacking. The purpose of this study was to validate a proof-of-concept method capable of quantifying 3D mitochondrial network morphology in whole mount skeletal muscle and then applying it to mitochondrial morphology analysis in cell types otherwise difficult to image within their native environment, terminal Schwann cells (tSCs). Herein, we report that mitochondrial networks were fragmented in dystrophic mouse muscle compared to healthy controls, as observed by others, and correlated with muscle pathology as expected. Using S100β reporter mice to identify Schwann cells, we labeled tSC mitochondrial networks *in vivo* prior to rapid imaging *in situ* with high-resolution confocal microscopy. Moreover, these methods offer a comprehensive and novel approach enabling the quantification of mitochondria network morphology across multiple cell types (like muscle fibers and tSCs) using standard microscopy available in university core facilities.

**Summary:** Local injections of mitochondrial dye are used to label terminal Schwann cells for confocal microscopy imaging after proof of concept was demonstrated in skeletal muscle tissue from mice with healthy or diseased muscle.

## Introduction

Mitochondria are remarkably plastic organelles acting as *the lungs of the cell*, consuming oxygen for cellular respiration to support adenosine triphosphate (ATP) production in response to metabolic demand ^1^. While once thought of as discrete organelles, it is now understood that mitochondria are highly integrated ^2–4^, sharing energy substrate throughout their network in accord with local requirements. Integration provides a distinct advantage over isolation, that is, connected mitochondrion support energy deficits maintaining electrochemical gradients for oxidative phosphorylation during challenge. Indeed, the distribution, volume, and morphology of mitochondria suit the energy needs in their vicinity ^5^. In health, regular turnover of mitochondrial networks is maintained through fusion and fission events ^2,3^. Fusion expands networks, promoting survivability and complementing deficits, while fission mitigates further mitochondrial damage through the preferential exclusion of dysfunctional regions ^2,3^.

Skeletal muscle comprises about 40% of body mass relying upon mitochondria for aerobic metabolism as well as cellular signaling to maintain homeostasis ^6^. Collectively, in conditions of skeletal muscle dysfunction, mitochondrial quality control mechanisms are impaired ^6^. Investigations of muscle dysfunction typically measure mitochondrial respiration, volume, morphology, and reactive oxygen species (ROS) production to assess mitochondrial quality ^3,6^. In that regard, during prolonged muscle dysfunction, mitochondrial respiration is reduced, while ROS are increased, and networks are fragmented ^3,6^. Perturbations in mitochondrial morphology arise largely from activation of mitochondrial fission proteins, dynamin related protein 1 (DRP1) and mitochondrial fission 1 (FIS1) ^2^. In response to muscle atrophy, DRP1 oligomerizes with multiple DRP1 dimers to create a filament around the mitochondria, decreasing in size until a given section of mitochondria are “pinched” off, promoting fragmentation ^3^. Together, over activity of mitochondrial fission paired with reductions in fusion of newly formed mitochondria are responsible for the fragmented appearance of dysfunctional mitochondrial networks, like those described in inactive or diseased muscle ^2^.

Mitochondrial morphology varies markedly between cell types, conforming to tissue function ^7^. Reports from liver tissues suggest that mitochondria are more compact and spherical while mitochondria in white matter of the brain are more elongated and tubular ^8^. Upon challenge as in conditions of aging and oxidative stress, mitochondria in these tissues exhibit a donut like phenotype. However, in osteoblasts from bone, mitochondrial donut formation and fragmentation were linked with increased mitochondrial secretion and osteoblast maturation ^9^. While skeletal muscle mitochondrial networks have been extensively examined in multiple species, limitations remain for the imaging of mitochondrial networks from certain cell types indwelling skeletal muscle, tSCs of the peripheral nervous system. These pose difficult to capture within their native environments as cell ultrastructure is significantly altered following isolation and these tissues are difficult to target with dye *in vivo* without also loading accompanying skeletal muscle.

Imaging techniques used to capture mitochondrial morphology in living systems can be categorized as either light or electron microscopy ^5^. With traditional light capture microscopy, dyes can label live cells, fixed cells can be immunolabeled, and mitochondria can be genetically labeled ^5^. Live cell imaging typically requires incubating living cells for 20-60 minutes prior to analyzing fluorescence. Dyes are directed to mitochondria via membrane potential and therefore may be altered under conditions of altered membrane potential ^5^. Capturing mitochondrial morphology of tSCs within skeletal muscle via dye loading is possible, however, it requires imaging of whole tissues and detailed labeling techniques. In particular, capturing mitochondria within intact structures without the interference of skeletal muscle labeling. Herein we report the use of a mitochondrial dye, MitoView Fix 640™, to label living skeletal muscle in health and disease prior to labeling tSC mitochondria *in vivo* and imaging *in situ*.

## METHODS

### Ethical approval

All protocols in this study were approved by Animal Care and Use Committee at Texas A&M University (AUP 2022-0215). Animal care was in accordance with the National Research Council’s Guide for the Care and Use of Laboratory Animals (Eighth Edition, 2011).

### Animals

Six-month-old (n=3) adult male DBA/2J [Wild-type (WT)] (RRID: IMSR_JAX:000671) and D2.B10-DMD*^D^*^2^*^.mdx^*/J [*D2.mdx* (mdx)] (RRID: IMSR_JAX:013141) mice were ordered from Jackson Labs (Bar Harbor, ME), while three to five-month-old (n=8) adult male and female B6;D2-Tg(S100-EGFP)^1Wjt^/J (S100B) (RRID:IMSR_JAX:005621) were used from an ongoing colony of investigator maintained mice (Figure 1).

**Figure 1.**
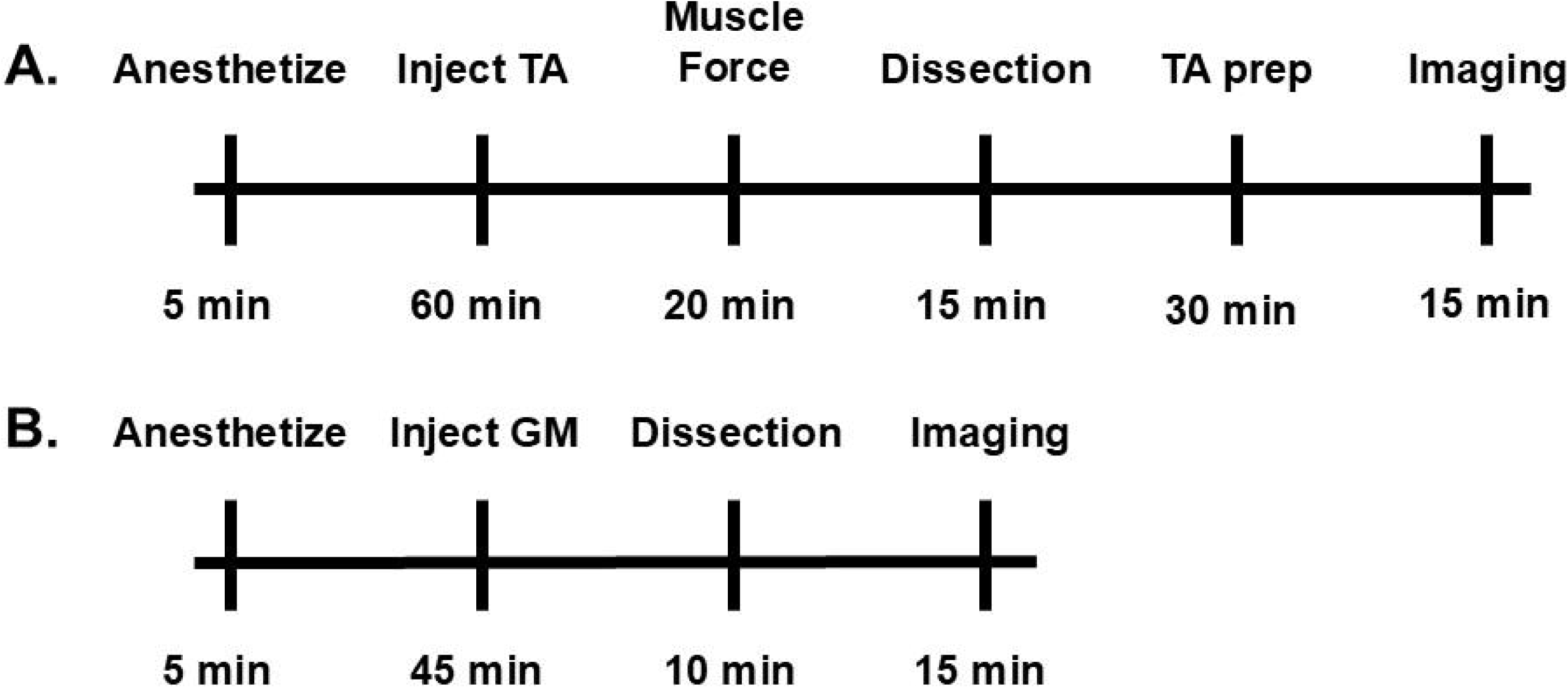
Laboratory workflow.

### In Situ Analysis of Peak Muscle Forces of an Intact TA Muscle

Following the 1-hour incubation period, mice were anesthetized using isoflurane (2–3% induction, 1.5–2% maintenance in oxygen). The left tibialis anterior (TA) muscle was surgically exposed while keeping the surrounding tissues intact to preserve physiological conditions as described ^10^, with the distal tendon of the TA muscle carefully tied to a load beam (1300A 3-in-1 Whole Animal System, Aurora Scientific, Ontario, Canada). A strip of KimWipe (Kimberly-Clark, Irving, TX, USA) was wrapped around the exposed muscle and irrigated (3 mL min–1) with 0.9% sterile saline, warmed to 40°C; a heat lamp maintained muscle temperature at 32°C. A pair of platinum/iridium wire electrodes were placed across the muscle belly for direct stimulation. The load beam was attached to a micrometer to adjust the optimal length (Lo) as determined during twitch contractions at 1Hz and recorded with a digital caliper. Maximum tetanic contractions were induced three times at 140 Hz with a 60 s rest between each contraction, with the highest force number recorded for measuring peak muscle force (g). Data was acquired using manufacturer software (610A Dynamic Muscle Control/Analysis, Aurora Scientific, Ontario, Canada) on a personal computer.

### Mitochondrial Dye Preparation, Loading, and Sample Treatment

- To optimize administration of mitochondrial dye *in vivo* for whole mount tissue staining preparations, 50 µl of a mitochondrial dye (Mitoview Fix 640, Biotium) suspended in sterile saline at a concentration of 2 µM was delivered via intramuscular injection to the right tibialis anterior (TA) muscle of anesthetized WT and *D2.mdx* mice as used in other TA injections ^11^.
- The dye incubated *in vivo* for 1 hour prior to experimentation to facilitate uptake in muscle mitochondrial networks prior to dissection.
- Isolated muscle fiber bundles from the excised TA under a stereo microscope (SZ61, Olympus, Breinigsville, PA, USA) were immediately fixed in 4% paraformaldehyde (PFA) at room temperature for 15–30 minutes during gentle teasing, stained with fluorescent Hoechst to label Nuclei, fluorescent α-bungarotoxin 552 (dilution 1:500; catalog. no. 00014; Biotrend, Koln, Germany) and washed in PBS three times for 5 min per wash.
- To visualize nuclear distribution, the fixed muscle fibers were incubated with Hoechst 33342 (Thermo Fisher Scientific) at a final concentration of 5–10 µg/mL in PBS for 15–20 minutes at room temperature.
- After staining, the samples underwent an additional three PBS washes (5 minutes each) to remove excess dye and reduce background fluorescence. For improved optical imaging, the muscle fiber bundles were treated with a commercially available tissue-clearing agent (Visikol). The clearing process followed the manufacturer’s instructions, with samples incubated in Visikol 1 and 2 for 30 minutes each at room temperature to achieve optimal tissue transparency.
- Confocal imaging was performed using a Stellaris 5 White Light Confocal Microscope (Leica Biosystems, Wetzlar, Germany) to obtain high-resolution three-dimensional (3D) optical sections of the muscle samples. Z-stack images were acquired at a depth of approximately 50 µm for muscle using a 63x magnification objective lens (N.A. 1.40).
- The imaging system was controlled using Leica LAS_X software (Leica Biosystems). While 640 nm is the reported excitation of Mitoview Fix, we discovered 646 nm to provide the greatest clarity with the least amount of background. Clarity was enhanced using Lightning Deconvolution (Leica Biosystems) mode, relying upon an advanced computational algorithm to refine signal-to-background ratios.
- To combat dehydration and photobleaching, tissues were transferred with saline, upon the cover slip side, and two drops of either Visikol 2

### tSC Mitochondrial Preparation

- S100B mice were used for imaging of tSCs.
- For tSC imaging, injections were made beneath the gluteus maximus muscle (GM) following a small skin incision as described by Fernando, C.A., *et al.*, (2018).
- Mitoview Fix 640, described above, was injected (75 µl) beneath the GM prior to wound closure.
- Following a 45-minute incubation time to label tSCs without significant labeling of muscle fibers, GMs were dissected, placed in chilled saline, and rapidly trimmed of excess fat before transferring to a cover slip for imaging.
- Confocal imaging was performed using a Stellaris 5 White Light Confocal Microscope (Leica Biosystems, Wetzlar, Germany) to obtain high-resolution three-dimensional (3D) optical sections of the muscle samples. Z-stack images were acquired at a depth of approximately 5-7 µm for tSCs using a 63x magnification objective lens (N.A. 1.40).
- The imaging system was controlled using Leica LAS_X software (Leica Biosystems). While 640 nm is the reported excitation of Mitoview Fix, we discovered 646 nm to provide the greatest clarity with the least amount of background. Clarity was enhanced using Lightning Deconvolution (Leica Biosystems) mode, relying upon an advanced computational algorithm to refine signal-to-background ratios.
- To combat dehydration and photobleaching, tissues were transferred with saline, upon the cover slip side, and two drops of Permount Mounting Medium (Cat. #SP15-100, Fisher Scientific, Waltham, MA 02454, USA) GM placed on the dorsal surface (20 µl) of GM prior to being flattened with a glass block (2.5 cm x 2 cm x 1 cm; mass 7.8 g).
- Focused ion beam scanning electron microscopy (FIB-SEM) was performed as described ^12^.

### Image Analysis

Quantitative analysis of confocal optical sections of TA was performed using AIVIA 3D machine-learning image analysis software (DRVision Technologies, Bellevue, WA, USA). Before automated segmentation, the software underwent multiple supervised training sessions to enhance algorithm adaptation. Following algorithm optimization, 3D image stacks were processed to quantify mitochondrial morphology, spatial distribution, and network connectivity. intensity thresholds, edge detection settings, and voxel size calibration were standardized across all samples to minimize variability.

For tSC, final optical sections were processed and analyzed using ImageJ/Fiji (NIH, Bethesda, MD, USA) and Leica LAS_X software. Quantitative assessments of mitochondrial morphology, network connectivity, and nuclear positioning were conducted using automated segmentation algorithms and threshold-based analyses. All imaging experiments were performed under identical conditions to maintain consistency and reproducibility across biological replicates.

### Immunohistochemistry

TA muscle samples were embedded in Tissue-Plus O.C.T. Compound (Scigen, Fischer Scientific, Hampton, NJ, USA) and sectioned (thickness, 10 μm) in a Cryostar NX50 Cryostat (Epredia, Kalamazoo, MI, USA) at -17°C onto a microscope slide. Muscle cross sections were stained as described^1^ to identify myofiber borders for cross-sectional area (CSA) analysis. The primary antibody used was rabbit anti-laminin (1:400, Cat.# NC1732938, Fisher Scientific; Hampton, NJ, USA), with secondary goat anti-rabbit Rhodamine (TRITC) (1:400, RRID: AB_90296, Cat.# AP132RMI), and slides mounted in Invitrogen ProLong Gold antifade reagent with DAPI (Cat.# P36941, Fisher Scientific, Hampton, NJ, USA). Slides were imaged on a Stellaris 5 White Light Laser confocal microscope (Leica Microsystems, Deer Park, IL, USA) using Leica LAS_X software (RRID: SCR_013673, Leica Microsystems, Deer Park, IL, USA). For each sample, 2-3 randomized 580 x 580 µm regions of interest were imaged with a 20x (N.A., 0.75) objective and values across all regions were averaged per muscle. Approximately 400 myofibers per TA muscle section were analyzed using semi-automatic muscle segmentation analysis in *NIS*-Elements Advanced Research software (RRID: SCR_014329, Nikon, Melville, NY, USA) presented as µm^2^.

### Western Blot Analysis

TA muscle samples from DBA.2J and *D2.mdx* mice were homogenized in lysis buffer (pH = 7.4) containing 5 mM Tris-HCI, 5 mM EDTA with protease and phosphatase inhibitors (Sigma-Aldrich, St. Louis, MO, USA). Homogenates were centrifuged at 10,000 *g* for 15 minutes at 4 °C. The supernatant (soluble fraction) was aspirated, and its protein concentration assessed using the Bradford method (Sigma-Aldrich, St. Louis, MO, USA). Protein samples (50 µg) were blotted and analyzed using Image Studio Lite as described^1^. Primary antibodies of interest were citrate synthetase (1:1000, RRID: AB_10678258) and OXPHOS Rodent Cocktail (1:500, RRID: AB_2629281). Membranes were exposed to IRDye 800CW Goat anti-Mouse IgG (1:20,000, RRID: AB_621842, Cat.# 926-32210) or IRDye 800CW Goat anti-Rabbit IgG secondary antibodies (1:20,000, RRID: AB_621843, Cat.# 926-32211) (Li-Cor Biotechnology, Lincoln, NE, USA) and scanned with a Li-Cor Odyssey DLx Imager (Li-Cor Biotechnology, Lincoln, NE, USA). Blots were normalized to total protein (A.U.) using Revert total protein stain (Cat.# 926-11011, Li-Cor Biotechnology, Lincoln, NE, USA). Individual data points are displayed as percentages corresponding to DBA.2J mean.

### Statistical Analysis

Summary data are reported as means ± S.E.M. Statistical analyses were performed using Prism 9 software (RRID: SCR_002798, GraphPad Software, La Jolla, CA, USA). Student’s t-tests were used to determine statistical significance among group mean differences. P ≤ 0.05 was considered statistically significant.

## RESULTS

### D2.mdx mice display phenotypic reductions in muscle function

To evaluate muscle pathology in diseased mice, muscle force was acquired via direct excitation. As reported previously^13^, muscle twitch (Tw) (g) and peak muscle force (P_0_) (g) produced by TA muscles of mdx mice was significantly less than WT group (means ± SE: Tw, WT 38 ± 4.426 g; mdx, 23 ± 15.33 g; P= 0.04; P_0_, WT 108 ± 11.44 g; mdx, 56.55 ± 3.3 g; P= 0.01) (Figure 2A and B).

**Figure 2.**
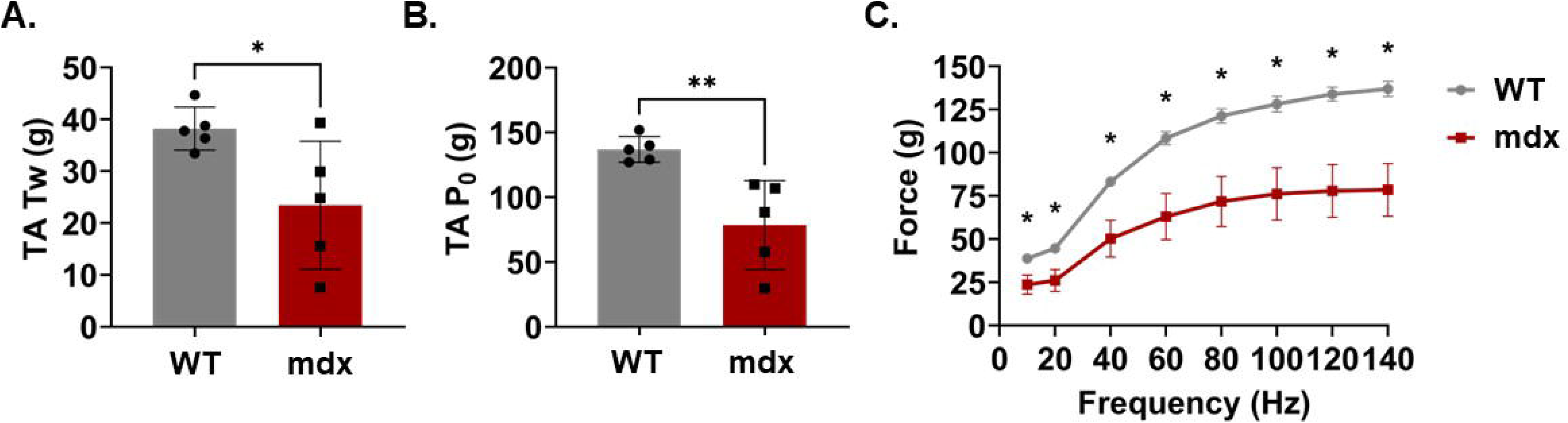
Comparisons of isometric force between WT and mdx mice. A) TA muscle twitch (Tw), B) peak muscle force (P_0_), and C) force frequency curve. Muscle functional parameters were reduced in mdx TA muscle compared to WT. Statistics: Students T-test. * indicates different from WT.

### Histology of mdx pathology

As a marker of histopathological characterization, myofiber CSA was binned into 400 μm^2^ increments to create a relative frequency histogram plot for WT and mdx groups. Consistent with reports the mdx phenotype shifts the CSA relative frequency curve to the left of WT with an almost 60% reduction in average fiber size (Figure 3).

**Figure 3.**
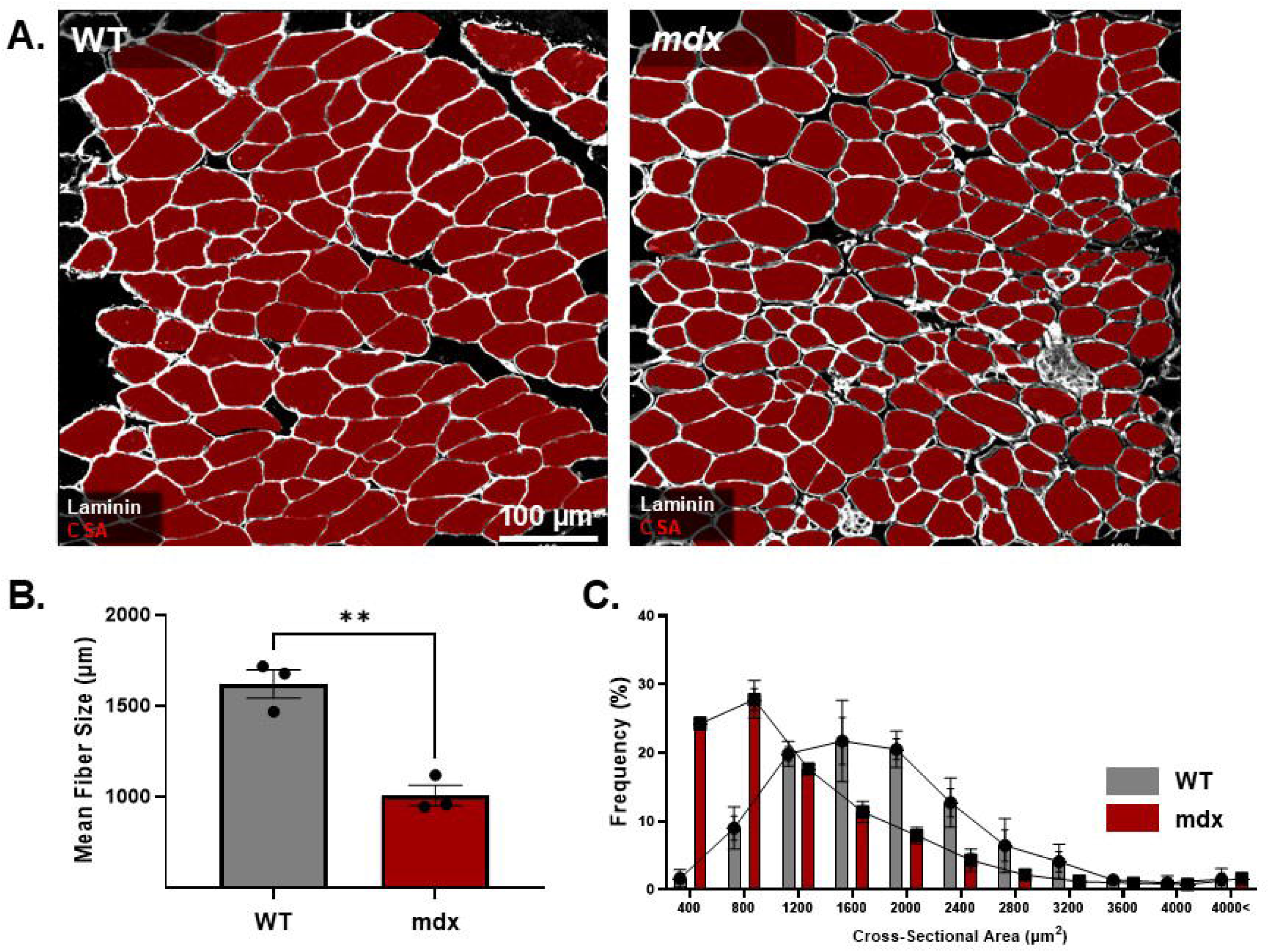
TA muscle CSA. A) Representative images of TA muscle cross sections with laminin (white) and myofiber interiors (red). Scale bar = 100 µm. B) Fiber size is significantly reduced and C) the fiber size frequency of occurrence was shifted to the left in mdx mice compared to WT. Statistics: Students T-test. * indicates different from WT.

### Mitochondrial networks displayed expected reductions in connectivity

The representative 3D optical images of TA muscle confocal scans in two types of group WT and mdx are presented in image, (Figure 4). In comparison with WT group, mitochondria connection appears reduced in mdx mice type (Figure 5A and B). Furthermore, mitochondria in dystrophy mice model appear fragmented, changing shape and morphology, which can be defined as abnormal mitochondria in dystrophy mice group. mdx mice exhibited an increase in small, fragmented mitochondrial networks [<100 µm^2^] (WT, 17.3 ± 3.5%; mdx, 36.3 ± 4.9%; P= 0.015) and a decrease in large interconnected mitochondrial networks [1000 µm^2^] compared to WT mice (WT, 23.5 ± 0.26%; mdx, 16.8 ± 1.02%; P= 0.001) (Figure 5B).

**Figure 4.**
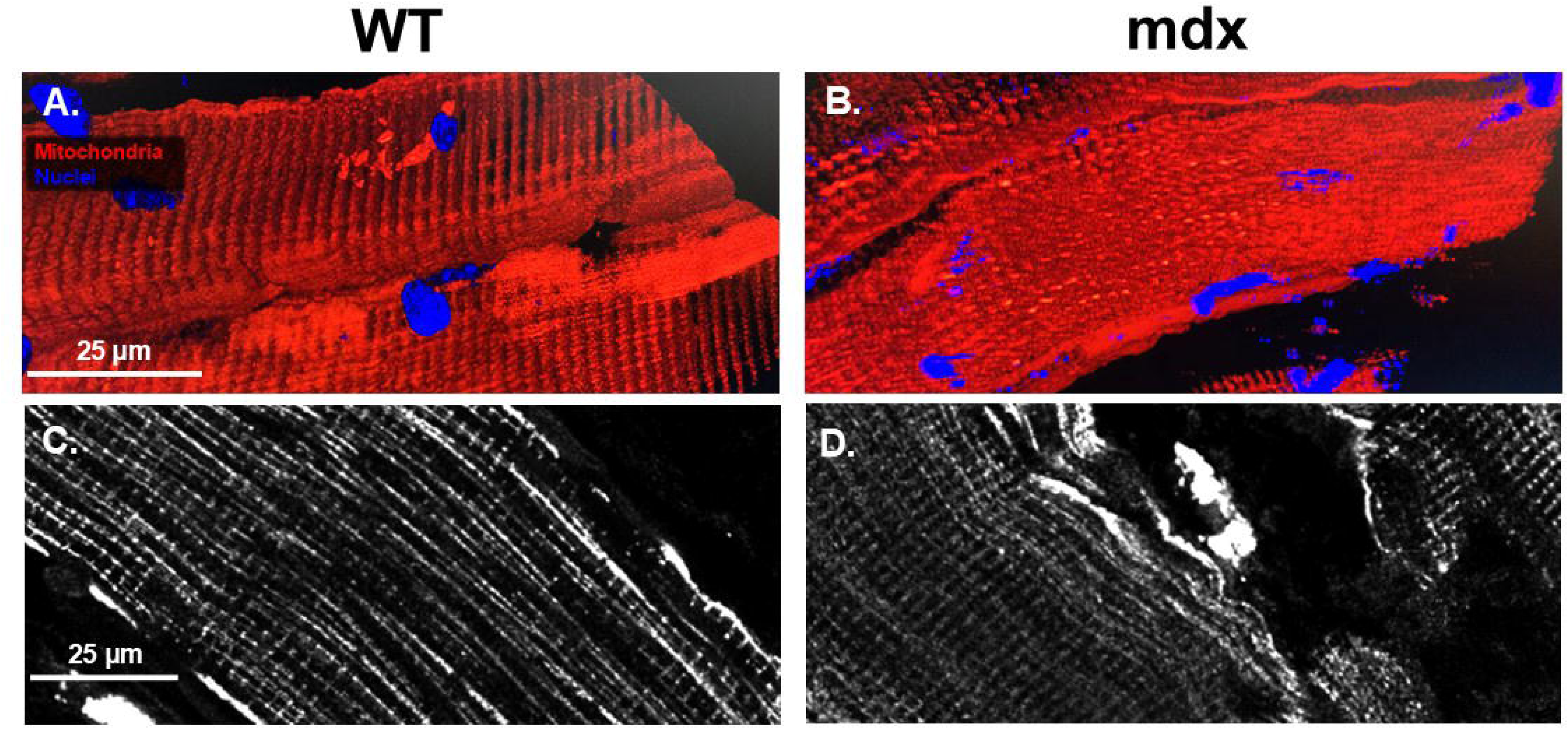
Representative images of TA muscle confocal scans from WT and mdx mice. A-B) 3D mitochondrial scans (red) with nuclei (blue) from WT and mdx mouse muscle. C-D) one micrometer thick optical sections of mitochondria (white) from healthy and dystrophic muscle. Scale bar = 25 µm.

**Figure 5.**
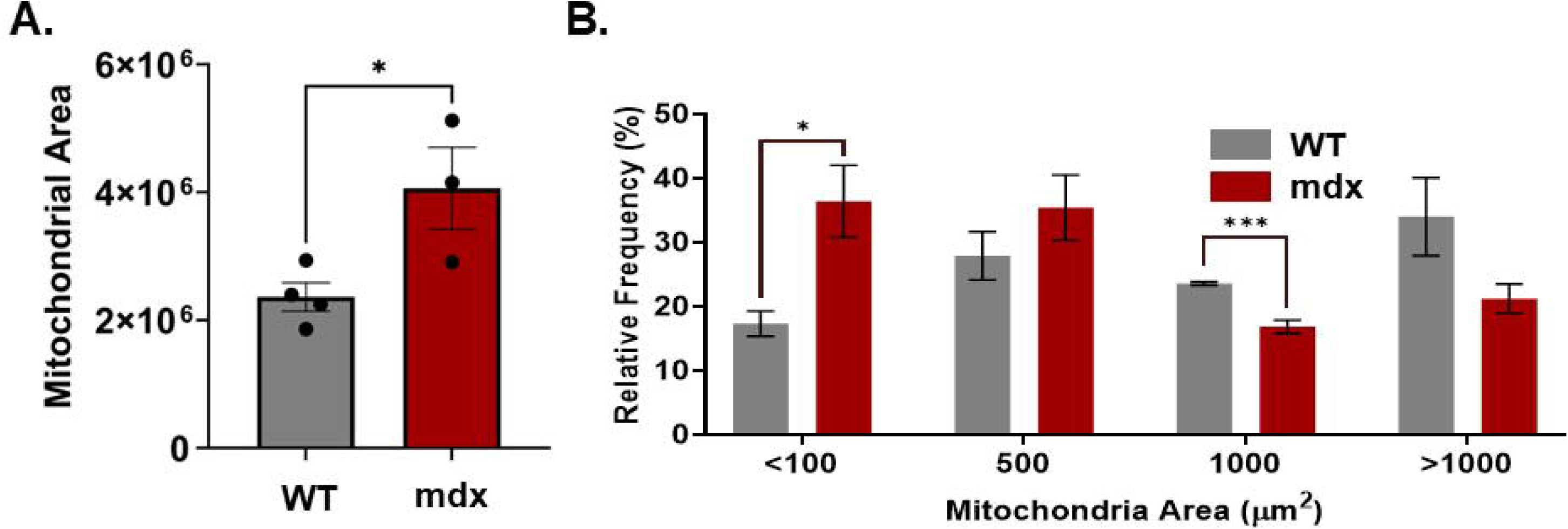
Quantification of mitochondrial morphological parameters in WT and mdx mouse TA muscle. A) Area of mitochondria measured throughout a Z-stack. B) Relative frequency of occurrence of discrete, mitochondria with areas binned by 500 µm^2^. Statistics: Students T-test. * indicates different from WT.

### Muscle function and histology correlates with mitochondrial connectivity

Linear regression was used to discern whether there was a correspondence between markers of muscle pathology and mitochondrial morphology. TA muscle mitochondrial networks with an area of 1000 µm^2^ or greater correlated to TA muscle force and CSA as indicators of D2.*mdx* muscle pathology (Figure 6A-H).

**Figure 6.**
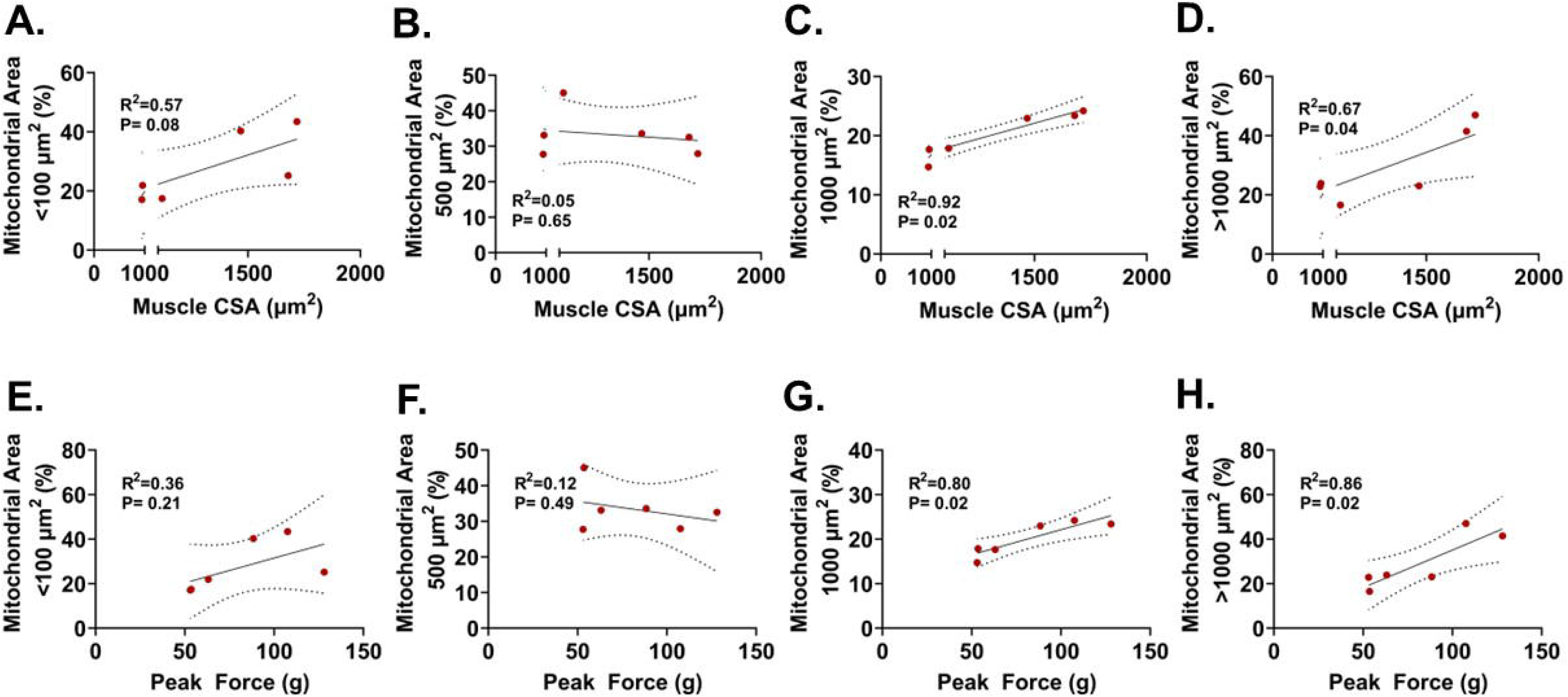
Analysis of mitochondrial morphology metrics relative to markers of mdx muscle dysfunction. For muscle CSA, frequency of occurrence of mitochondrial areas of 1000 µm^2^ or greater correlated (C and D), with a trend observed in mitochondrial areas of less than 100 µm^2^ (A), and no trends with mitochondria at 500 µm^2^. Muscle force correlated with only mitochondrial areas of 1000 µm^2^ or greater (G and H), only slight trends were observed for mitochondrial areas of less than 100 and 500 µm^2^ (E and F). Statistics: linear regression with Pearson’s Correlation determined slope deviation from zero.

**Figure 7.**
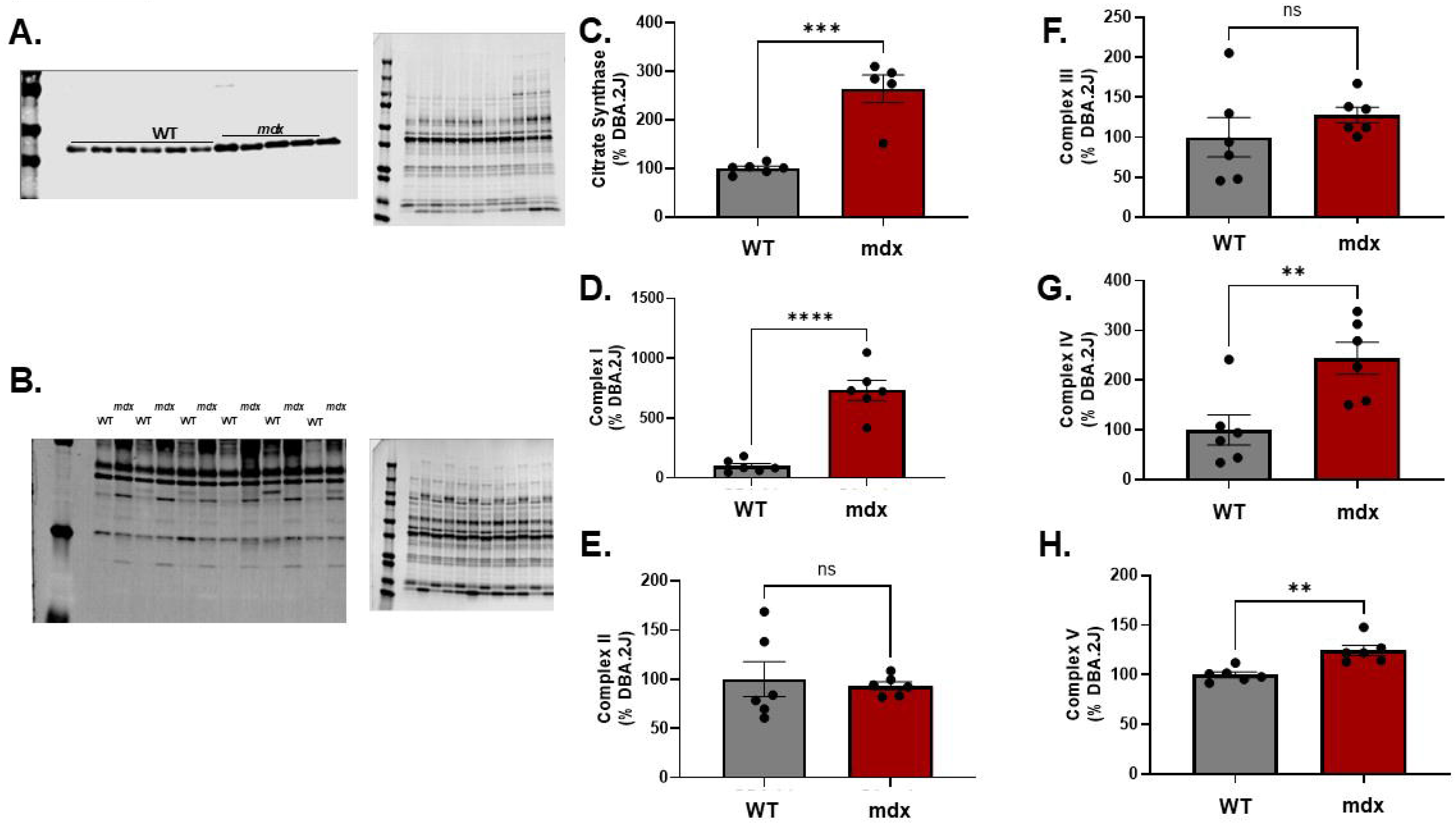
Citrate Synthase and oxidative phosphorylation complexes are increased in mdx compared to WT. A) Representative immunoblots and B) total protein for mitochondrial protein quantification. C) Citrate Synthase protein abundance. D-H) Oxidative phosphorylation complexes I-V. Blots were normalized to total protein stain. Statistics: Students T-test. * indicates different from WT.

### Mitochondrial Proteins are increased

Given that mitophagy is reduced in muscle dystrophy^14^ and confocal imaging revealed increased fragmentation and volume of mitochondria, we evaluated protein abundance of mitochondrial proteins. Citrate Synthase was used as a surrogate marker of mitochondrial content and significantly increased ∼150% in mdx over WT. Regarding mitochondrial complexes, complexes I, IV, and V were significantly elevated in mdx TA muscle over WT.

### tSC imaging in situ

Fluorescent confocal microscopy revealed mitochondrial morphology of tSCs in situ. Both tSC morphology and mitochondrial morphology are preserved following labeling and rapid removal, sample preparation, and imaging. Mitochondria of tSCs were observed to have interconnected loops (Figure 8A, C-D) near bifurcations. Image analysis revealed that (red) mitochondria make up <30% of tSC (green) (Figure 8E-G).

**Figure 8.**
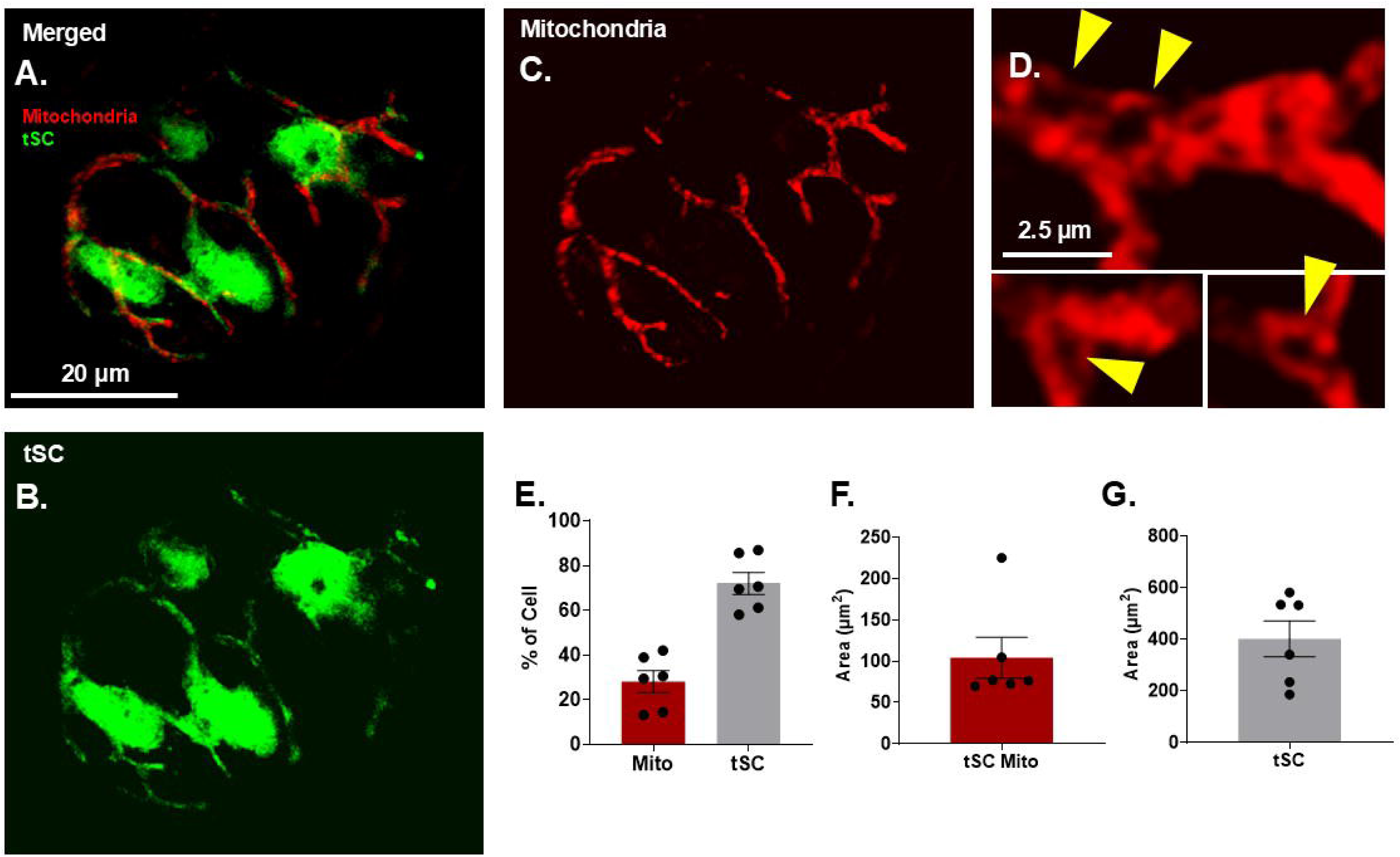
tSC mitochondrial morphology. A-C) tSC in green and mitochondria in red. Scale bar = 20 µm. D) Close images of mitochondrial bifurcations, yellow arrows depict interconnected loops of mitochondria at each bifurcation. Scale bar = 2.5 µm. E-G) Quantified parameters of cell area occupied by mitochondria.

Further validation of tSC mitochondrial morphology was performed qualitatively comparing fluorescent confocal imaging to focused ion beam scanning electron microscopy demonstrating similar doughnut shapes in tSC mitochondria (Figure 9A-D). Analysis of post synaptic components revealed subsarcolemmal mitochondria, void of the characteristic pretzel like configuration (as in Figure 9E), but clustered around AChRs embedded upon the surface of muscle fibers (Figure 9F-H).

**Figure 9.**
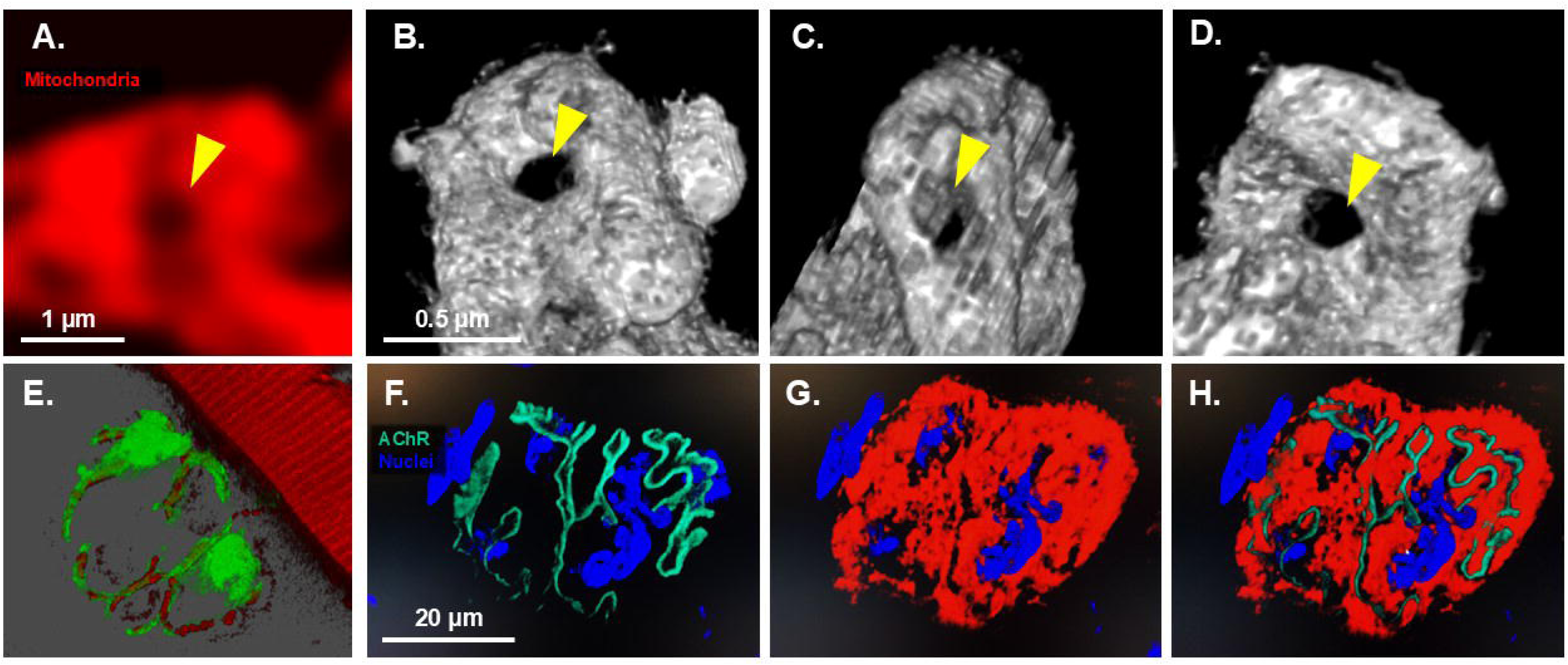
Validation of tSC mitochondrial imaging. A) zoom of mitochondrial doughnut (yellow arrow) with confocal microscopy. Scale bar = 1 µm. B-D) images of tSC mitochondrial doughnut at multiple angles using focused ion beam scanning electron microscopy. Scale bar = 0.5 µm. E) 3D image of green tSC with red mitochondria and adjacent red mitochondria in neighboring muscle fiber. F) 3D image of acetyl choline receptors (AChR) (teal) and nuclei (blue), G) sub sarcolemma mitochondria (red), and nuclei, H) merged. Scale bar = 20 µm.

## DISCUSSION

There are four primary tissue types in the human body: epithelial, connective, muscle, and nervous tissue. Each of these tissue types are unique in structure and function, reflected in energy demand and morphology. Mitochondrial location within cellular structures as well as morphology correlates strongly with mitochondrial health and metabolic demand. Given the clinical relevance of mitochondrial health coupled with divergent respiration kinetics as well as morphologies of different cell types, we sought to develop an easily applied, confocal specific method for evaluation of mitochondrial morphologies in multiple cell types, providing information on mitochondrial health as indicated by morphology, particularly in tissue types difficult to maintain *in vivo* structure upon isolation.

### Mitochondrial morphology in healthy and diseased skeletal muscle

Central to the development of a novel technique, the experimentation of the new technique was implemented against a validated control. Skeletal muscle is 40% of body mass and necessary for breathing, myokine secretion, circulating factor uptake, and locomotion. Owing to its high body mass, ease of isolation, and multiplicity of physiological effects, mitochondrial morphology has been investigated in skeletal muscle since the 1960s ^15^.

Mitochondrial morphology in skeletal muscle has been closely associated with both health and disease ^2,7,15^. Indeed, in genetic muscle diseases like muscular dystrophies, mitochondria are shown to be more fragmented, generating greater amounts of reactive oxygen species (ROS), reduced metabolic efficiency, and contributing to a greater overall pathology ^16^. Conversely, focusing on restoration of healthy mitochondrial networks and morphology in disease results in improvements in markers of myopathy, that is, improved contractile performance, reduced fibrosis, and lower net production of ROS ^14^.

Consistent with reports of skeletal muscle mitochondrial morphology in the context of mouse models of Duchenne Muscle Dystrophy, our method of directly injecting skeletal muscle with dye attracted to the negative membrane potential of mitochondria revealed fragmented mitochondria and increased expression of surrogate protein markers of mitochondrial volume. Additionally, these markers of mitochondrial pathology were also highly correlated with traditional markers of muscle pathology, specifically, muscle function and cross-sectional area.

### tSC Mitochondria

Schwann cells provide insulation to axons for rapid transmission of action potentials ^17^. Given their importance, Schwann cell disruption causes both sensory and movement disorders like Multiple Sclerosis and Charcot-Marie-Tooth disease. Notably, mitochondrial diseases often result in neuropathic phenotypes while defects in peripheral nervous system axons often correlate with damage to mitochondrial DNA ^18^. Following investigations of mitochondrial abnormalities using electron microscopy in cross sections of peripheral nerves, it has been noted that mitochondria appear swollen and malformed, primarily in the enshrouding Schwann cells versus axon ^19,20^. While poorly understood, there is a strong association between peripheral Schwann cell mitochondria morphology and neuropathology.

Recent evidence suggests that tSCs are particularly important in regard to diseases like spinal muscular atrophy and Amyotrophic lateral sclerosis ^21^. Remek Schwann cells or tSCs are unique in peripheral glia as they remain unmyelinated, enshrouding the neurofilaments hovering above synaptic clefts of the neuromuscular junction (NMJ) ^21,22^. The NMJ is the active site of neuromotor activity. In particular, axonal propagation down the axon promotes a release of acetylcholine to activate myofibers for contraction. Following injury, tSCs aid in the recovery of NMJs. Concomitant with malformed NMJs in neuromotor pathologies, tSC morphology is equally malformed.

Despite the close association of mitochondrial health and morphology to pathology, little to no work has been conducted on whole mount imaging of tSC mitochondria with light microscopy. Given that laser scanning confocal microscopy is readily available to most university research cores, we determined to develop a method to image and quantify mitochondrial morphology in tSCs *in situ*. Notably, the most striking characteristics of our analysis of healthy tSC mitochondria are the loops found at each bifurcation of the recognizable pretzel like configuration (Figure 8 C and D and 9 A-D). While stand-alone loops and individual mitochondrial globules are associated with mitochondrial fission, interconnected segments and loops are indicators of fusion events ^23^, culminating in healthy mitochondrial interconnectivity and participation in substrate shuttling.

## Conclusions

Mitochondrial morphology has been used to draw implications around mitochondrial health and locations of metabolic activity. Herein we show that injecting mitochondrial dye into soft tissues in vivo provides unique advantages for understanding mitochondrial morphology in tissues otherwise difficult to obtain, i.e., tSCs. The present findings demonstrate a highly interconnected network of mitochondria within each tissue type. Disconnected mitochondria correlate with markers of dysfunction, consistent with other reports, demonstrating the validity of this approach. We conclude that this technique is useful for low abundance tissues arranged within a specific morphological configuration that is typically destroyed upon dissection.

## DATA AVAILABILITY STATEMENT

The data that support the findings of this study are available from the corresponding author upon reasonable request.

## AUTHOR CONTRIBUTIONS

Conceptualization: A.B.M., J.A.K., Data Curation: A.B.M., J.A.K., Formal Analysis: A.B.M., J.A.K., S.G., Funding Acquisition: A.B.M., J.A.K., Investigation: A.B.M., J.A.K., S.G., A.G.N., Methodology: A.B.M., J.A.K., B.G., A.G.N., Project Administration: A.B.M., Resources: A.B.M., J.A.K., Validation: A.B.M., J.A.K., Visualization: A.B.M., J.A.K., B.G., Writing – original draft: A.B.M., S.G., Writing – review and editing: A.B.M., J.A.K., S.G., B.G., A.G.N., All authors have read and approved the final version of this manuscript and agree to be accountable for all aspects of the work ensuring that questions related to accuracy or integrity of any part of the work are appropriately investigated and resolved. All persons designated as authors qualify for authorship and all those who qualify for authorship are listed.

## FUNDING

Supported by the Sydney and J.L. Huffines Institute for Sports Medicine and Human Performance Student Research Grant (JAK) and funds from the College of Education and Human Development at Texas A&M University (ABM).

